# Higher prevalence of *Bacteroides fragilis* in Crohn’s disease exacerbations and strain-dependent increase of epithelial resistance

**DOI:** 10.1101/2020.05.16.099358

**Authors:** Heike E. F. Becker, Casper Jamin, Liene Bervoets, Pan Xu, Marie J. Pierik, Frank R. M. Stassen, Paul H. M. Savelkoul, John Penders, Daisy M. A. E. Jonkers

**Affiliations:** Department of Medical Microbiology, NUTRIM School of Nutrition and Translational Research in Metabolism, Maastricht University Medical Centre+, Maastricht, The Netherlands; Division of Gastroenterology/Hepatology, Department of Internal Medicine, NUTRIM School of Nutrition and Translational Research in Metabolism, Maastricht University Medical Centre+, Maastricht, The Netherlands; Department of Medical Microbiology, Caphri School for Public Health and Primary Care, Maastricht University Medical Centre+, Maastricht, The Netherlands; Department of Medical Microbiology and Infection Control, Amsterdam University Medical Center, location VUMC, Amsterdam, The Netherlands

**Keywords:** *Bacteroides fragilis*, Crohn’s disease, barrier function, prevalence, metabolomics, whole genome sequencing, organoids

## Abstract

*Bacteroides fragilis* has previously been linked to Crohn’s disease (CD) exacerbations, but results are inconsistent and underlying mechanisms unknown. This study investigates the epidemiology of *B. fragilis* and its virulence factors *bft (*enterotoxin) and *ubiquitin* among 181 CD patients and the impact on the intestinal epithelial barrier *in vitro*.

The prevalence of *B. fragilis* was significantly higher in active (n=69/88, 78.4%) as compared to remissive (n=58/93, 62.4%, p=0.018) CD patients. Moreover, *B. fragilis* was associated with intestinal strictures. Interestingly, the intestinal barrier function, as examined by transepithelial electrical resistance (TEER) measurements of Caco-2 monolayers, improved when exposed to secretomes of *bft*-positive (increased TEER ∼160%, *p*<0.001) but not when exposed to *bft-* negative strains. Whole metagenome sequencing and metabolomics, respectively, identified 19 coding sequences and two metabolites that discriminated TEER-increasing from non-TEER-increasing strains.

This study revealed a higher *B. fragilis* prevalence during exacerbation. Surprisingly, *bft-*positive secretomes improved epithelial resistance.

## Introduction

Crohn’s disease (CD) is a chronic inflammatory disease, characterized by patchy inflammation of the intestinal mucosa with or without extra-intestinal manifestations.^1^ The disease course varies largely among patients with alternating periods of remission and exacerbations. Insufficient control of the recurrent mucosal inflammation results in phenotype progression and complications, such as strictures of fistulas, contributing to a high disease and economic burden.^2,3^

CD onset is considered to involve genetic predisposition, environmental factors and an adverse immune reaction to the host microbiota.^1^ However, the factors influencing the occurrence of exacerbations, complications and disease phenotype remain largely unclear. In recent years, microbial dysbiosis gained increasing attention as a factor contributing to exacerbations.^1,4^ Several studies reported a decreased microbial diversity and altered microbial composition in active CD patients compared to remission.^4–6^ As a consequence of compositional changes, alterations in overall microbial functionality are conceivable.^7^ Since the intestinal epithelium limits bacterial attachment by a mucus layer, bacteria often interact with the host via their secretome, consisting of metabolites, proteins and bacterial membrane vesicles (MVs).^8–11^ On the one hand, secreted glycosidases and mucinases are able to degrade the mucus layer and allow microbes to directly interact with epithelial cells.^10,12^ On the other hand, bacterial metabolites, such as the short chain fatty acid butyrate, have been reported to promote mucus production and the sealing capacity of the intercellular junctional complex.^13,14^ In CD, an impaired intestinal epithelial barrier function has increasingly been recognized as a hallmark of exacerbations.^1^ The impaired barrier function has been associated with alterations in the epithelial junctional complex.^15,16^ It remains unknown whether the observed altered microbiota composition and functionality during CD exacerbations can contribute to this impaired epithelial barrier.

In addition to alterations in microbial diversity, specific microbial taxa were detected in CD exacerbations^4,17,18^ of which *Bacteroides fragilis* is a prominent example.^4,17,19^ Several studies investigated the colonization of *B. fragilis* during exacerbation and remission in fecal samples and biopsies. Together, the results are inconclusive and based on rather low sample sizes.^4,17,19,20^ So far, it is still unclear whether the prevalence of *B. fragilis* differs in different disease stages of CD and how it may affect disease activity.

*B. fragilis* is a Gram-negative commensal of the phylum Bacteroidetes and some strains secrete various virulence factors, such as an eukaryotic-like ubiquitin (Ubb) and *B. fragilis* toxin (Bft; fragilysin).^21,22^ It is yet unclear whether these virulence factors contribute to CD exacerbations.

To gain more insights into the interaction between the intestinal microbiota and the intestinal barrier in CD, especially during dysbiosis, research essentially needs to focus on microbial functionality, such as reflected by the secretome. In this study, we therefore aim to investigate firstly the prevalence and relative abundance of *B. fragilis* and virulence-factor positive strains in a large, well-defined cross-sectional CD patient cohort and secondly the impact of the *B. fragilis* secretome on the intestinal epithelial barrier as first site of interaction and pathophysiological factor in CD exacerbations. We hypothesize that *B. fragilis* and its virulence factors *bft* and *ubb* are involved in exacerbations and by disrupting the intestinal epithelial barrier.

## Results

### Patient population

In total, 181 CD patients, 88 with active disease and 93 in remission, from the IBD South Limburg (IBDSL)^23^ cohort fulfilling the inclusion criteria were available for the present study. Baseline characteristics were comparable between patient groups, although steroid use was slightly higher among patients with active disease (table 1).

**Table 1.** Baseline characteristics of CD patient cohort.

|  | CD remission (n=93) | CD exacerbation (n=88) |
| --- | --- | --- |
| **Age** (median in years; IQR) | 43.0 (32.0-56.5) | 45.0 (28.0-57.0) |
| **Gender** (% female) | 63.4 | 51.1 |
| **Calprotectin** (median; IQR) | 22.0 (14.0-55.5) | 476.5 (325.8-703.0) |
| **Montreal classification** (%) |  |  |
| A1; A2; A3 | 6.5; 67.7; 25.8 | 10.2; 61.4; 28.4 |
| L1; L2; L3; L4 | 35.5; 17.2; 47.3; 4.3 | 23.9; 21.6; 54.5; 11.4 |
| B1; B2; B3; p | 59.1; 19.4; 31.2; 17.2 | 62.5; 25.0; 17.1; 11.4 |
| **Surgery** | 34.4 | 31.8 |
| **IBD medication** use (%) |  |  |
| Thiopurine | 34.4 | 29.6 |
| Biologicals | 58.1 | 48.9 |
| Steroids | 17.2 | 30.7 |
| Other | 13.4 | 18.2 |

L1 = Terminal ileum; L2 = Colon; L3 = Ileocolon; A1 < 16 years, A2 = 17-40 years, A3 > 40 years at diagnosis; B1 = without structure formation, non-penetrating; B2 = Stricturing; B3 = Penetrating

### *B. fragilis* prevalence and relative abundance

Based on the detection of the *B. fragilis* specific *gyrB* gene, the prevalence of *B. fragilis* was significantly higher among CD patients during an exacerbation (78.4 %) when compared to those in remission (62.4 %, *p* = 0.018, figure 1A). We subsequently determined the prevalence of the *B. fragilis* virulence factors *bft* and *ubb*. Only 11.4 % and 7.5 % (*p* = 0.376) of the samples from the exacerbation and remission group, respectively, were positive for *bft*. When considering only samples that were tested positive for *B. fragilis*, this equals to a proportion of *bft*-positive samples of 14.5 % and 12.1 % (*p* = 0.689), respectively. For *ubb*, the prevalence was 19.3 % and 11.8 % (*p* = 0.164) in the entire exacerbation and remission group, respectively, and 24.6 % and 19.0 % (*p* = 0.442), when considering the *gyrB* positive samples only (figure 1A). The relative abundance of *gyrB, bft* and *ubb* was not significantly different between remission and exacerbation (figure 1B-D).

**Figure 1.**
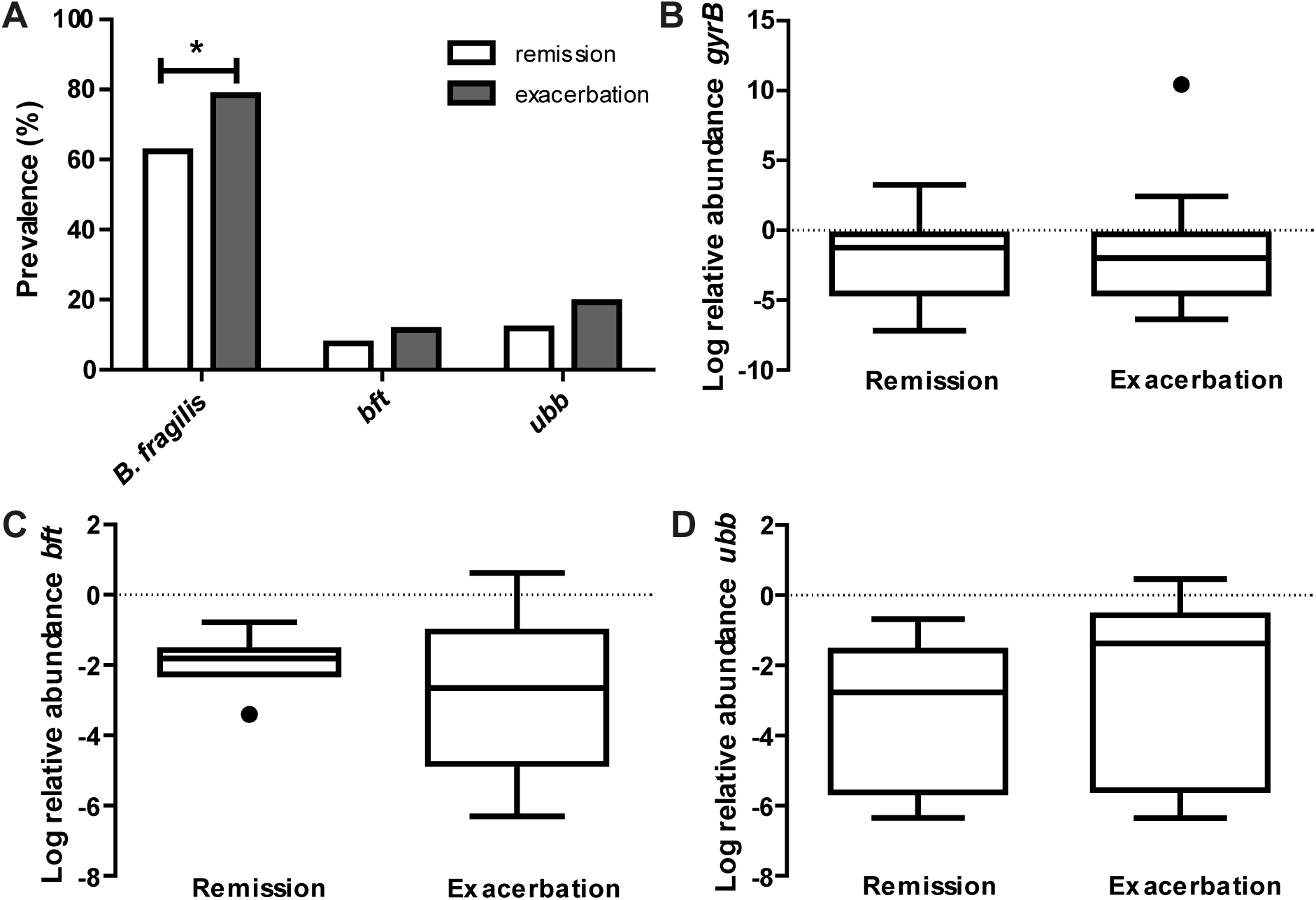
*B. fragilis* is more prevalent in CD exacerbation samples. (A) The prevalence of *B. fragilis* specific *gyrB* is 16.0 % higher in active CD compared to remission (\**p* = 0.018), while the prevalence of the virulence factors *bft* and *ubb* does not differ significantly between groups (*p* = 0.376 and *p* = 0.164, respectively). The relative abundance of *gyrB* (B), *bft* (C) and *ubb* (D), based on 16S rRNA gene copy number, does not significantly differ between active CD and remission (*p* = 0.837, *p* = 0.283 and *p* = 0.196, respectively).

Based on multivariable logistic regression, we found the following patient characteristics to be correlated with overall *B. fragilis* colonization: a stricturing disease behavior (Odds ratio (OR) = 3.212; *p* = 0.018), active Crohn’s disease (OR = 2.048; *p* = 0.038), and previous intestinal resections (OR = 0.406; *p* = 0.016).

### *B. fragilis* strain isolation and characteristics

In order to examine the impact of *B. fragilis* and its virulence factors on barrier function, we selected five *B. fragilis* strains cultured from samples of two healthy controls (HC-1, HC-2) and three CD patients (CD-1, CD-2, CD-3) and additionally included one reference strain (ATCC^®^ 25285(tm)). Based on whole genome sequencing, pairwise genetic distance revealed only 160 SNPs between the CD-2 (*bft-*positive) and the CD-1 (*bft-*negative) strains, whereas the median distance was 16960 SNPs among all strains (figure 2A). The genomic similarity was 89.7 %, based on shared split kmers between these two strains using Jaccard dissimilarity (mean Jaccard index 59.9 %). This indicates a large shared genomic backbone in these two strains. Sanger sequencing confirmed the presence of *bft-1* in HC-1, CD-2 and CD-3 and the presence of *ubb* in HC-1 and ATCC^®^ 25285(tm) with 93-99 % and 96-99 % sequence similarity, respectively. HC-2, ATCC^®^ 25285(tm) and CD-1 were confirmed negative for *bft*, and HC-2, CD-1, CD-2 and CD-3 for *ubb*. Cell rounding of subconfluent HT-29 cells could be detected in response to the supernatants of all bft-positive strains, which was interpreted as Bft activity (figure 2B). However, the concentration of Bft was found to be below the detection limit of western blot analysis (figure 2C). In addition, no cell rounding was detected when applying concentrated culture supernatant of *bft-1* positive strains on Caco-2 cells (data not shown) or colonic organoids (figure 3D), which suggests a cell type-dependent response to Bft. No cell rounding was detected on *bft-1* positive strains derived MVs, suggesting that the isolated MVs do not contain or release relevant amounts of (active) Bft (figure 2D).

**Figure 2.**
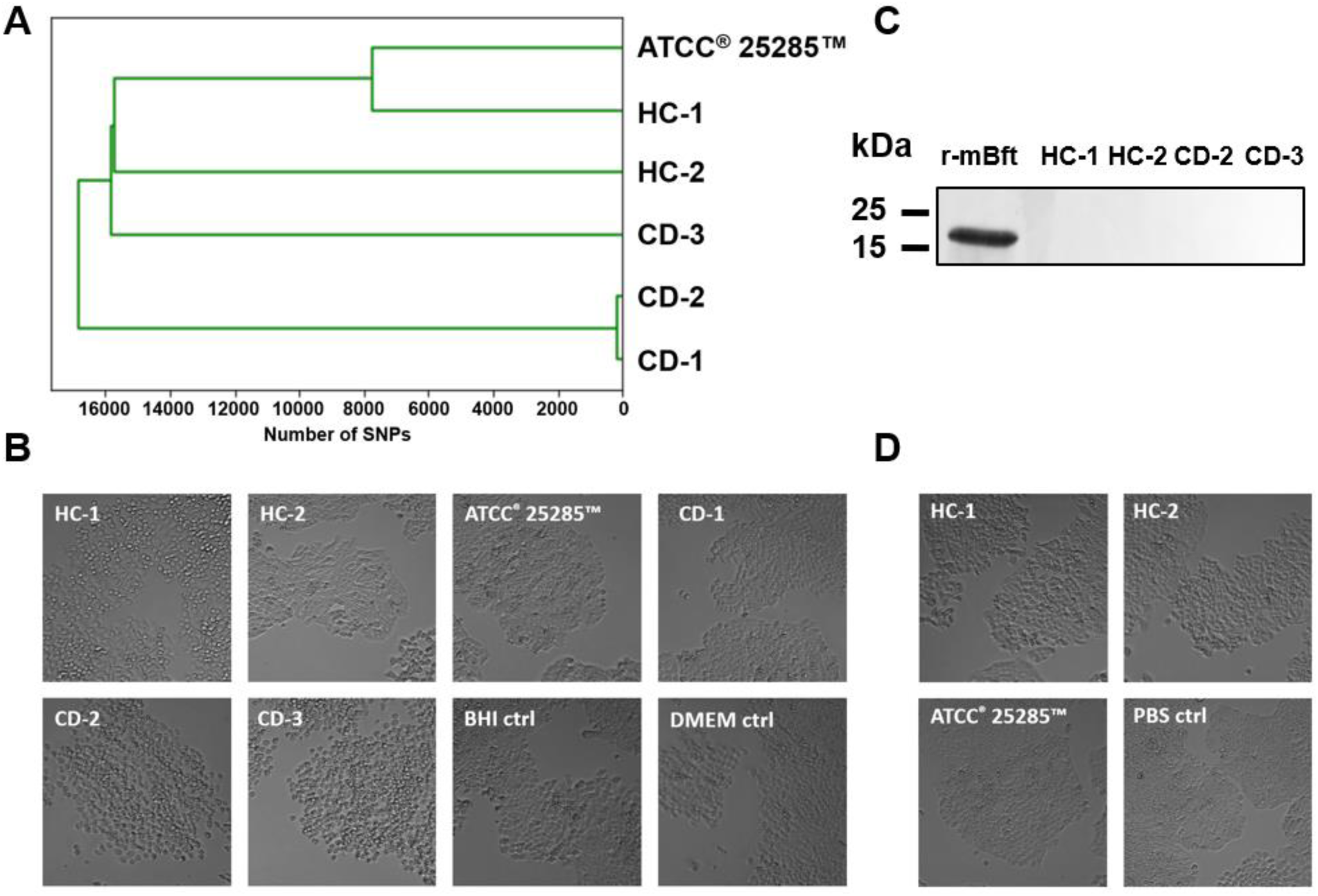
Characteristics of cultured *B. fragilis* strains. (A) Based on whole genome sequencing, pairwise genetic distance revealed 160 SNPs between CD-1 and CD-2, whereas the median distance was 16960 SNPs. (B) Cell rounding of subconfluent HT-29 cells could be detected in the supernatants of all *bft*-positive strains (HC-1, CD-2, CD-3). (B) Mature Bft could not be detected in the supernatant of *bft*-positive strains when compared to 200 ng of recombinant mature Bft (r-mBFT). (D) Cell rounding of subconfluent HT-29 cells could not be detected in response to *bft-*positive strain-derived MVs.

**Figure 3.**
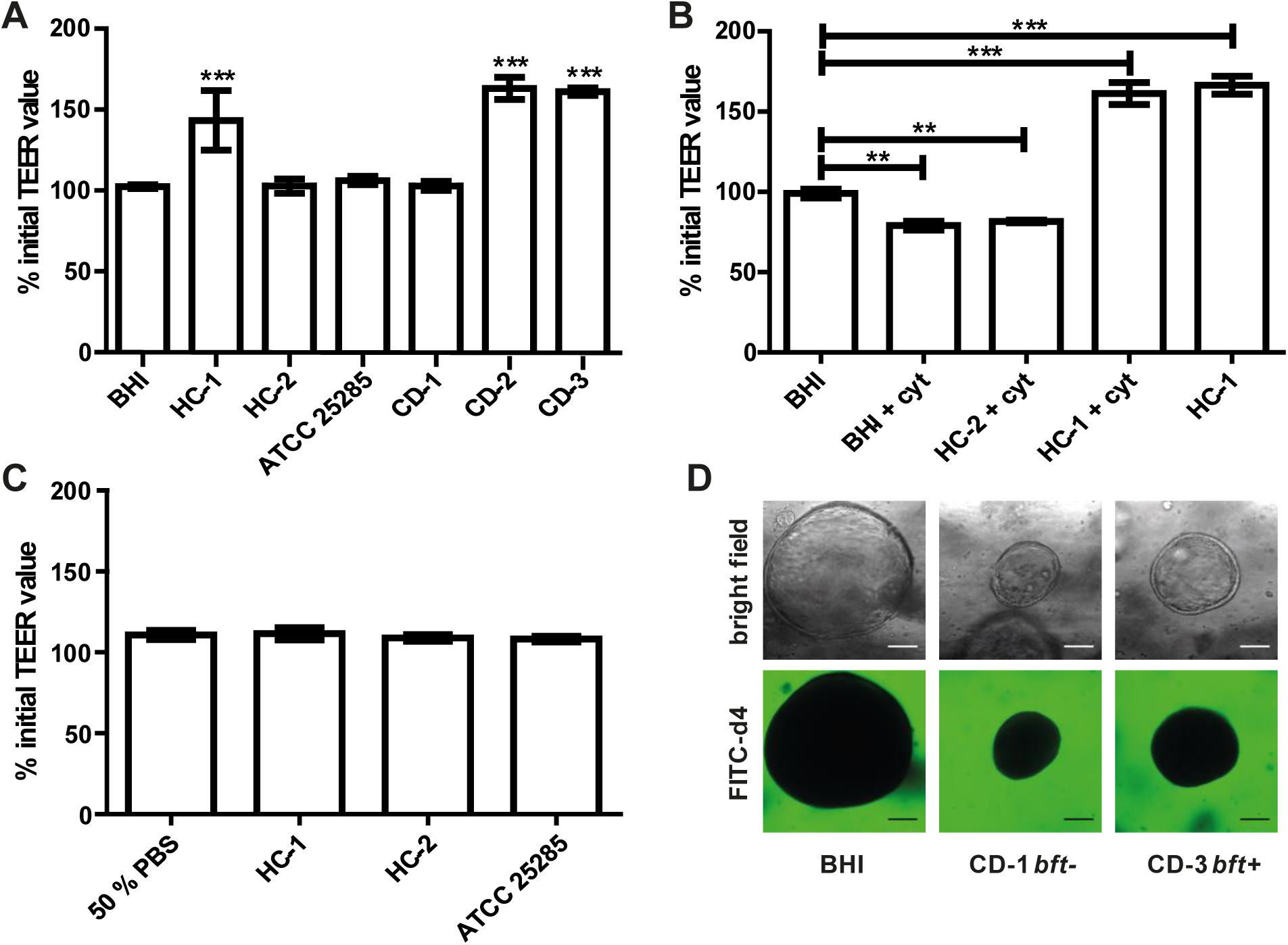
Barrier function examination of Caco-2 monolayer and colonic organoids exposed to *B. fragilis* culture supernatant and MVs. (A) Luminally applied *B. fragilis* concentrated culture supernatant of HC-1, CD-2 and CD-3 led to increased TEER after for 24 h. (B) Pre-incubation with 100 ng/ml TNF-α and IFN-γ could not impair the TEER enhancing effect of HC-1. (C) Luminally applied MVs, isolated from different *B. fragilis* strains, did not lead to TEER alterations in Caco-2 cell monolayers after 24 h. (D) CD patient-derived colonic organoids treated basally with *B. fragilis* CD-1 and CD-3 concentrated supernatant and FITC d4 (green) remain a stable barrier after 24 h incubation. Means +/-SD was based on triplicates of one representative experiment. cyt = cytokines; BHI = brain heart infusion broth

### Barrier modulation by *B. fragilis* supernatant and MVs

After 24 h exposure to concentrated supernatant of *bft-*positive strains, no decrease in transepithelial electrical resistance (TEER) was detected. Instead, TEER values increased up to 160 % (*p* < 0.001) when compared to brain heart infusion broth (BHI) control and to the concentrated culture supernatant of *bft-*negative strains (figure 3A). A similar increase by *bft-* positive supernatant was also observed after 1 h basal pre-incubation with 100 ng/ml tumor necrosis factor α (TNF-α) and 100 ng/ml interferon γ (IFN-γ), while *bft-*negative culture supernatant did not alter TEER (figure 3B).

Besides culture supernatant, we also examined the effect of MVs, which are vesicles released by Gram positive and Gram negative bacteria and can deliver a variety of products, including toxins, to host cells or neighboring bacteria.^8,11,24^ After 24 h incubation with 5 × 10^7^ MVs/ml, no alterations in TEER were observed in differentiated Caco-2 cell monolayers (figure 3C). Using a more physiological patient-derived colonic organoid model and a functional fluorescein isothothiocyanate-labeled dextran of 4 kDa (FITC-d4) permeation assay, we confirmed that the supernatant of different *B. fragilis* strains did not disrupt the epithelial barrier nor altered the morphology upon 24 h basal incubation (figure 3D).

### Junctional alterations in response to *B. fragilis* supernatant

To further elucidate the mechanisms underlying the TEER enhancing effect of Bft-positive culture supernatant, we performed qPCR to analyze the mRNA expression of various junctional genes in Caco-2 cells. When comparing one *bft-*positive (CD-3) and one *bft-*negative (CD-1) strain in quadruplicate, we could not detect any significant alterations in gene expression levels of junctional genes (figure 4).

**Figure 4.**
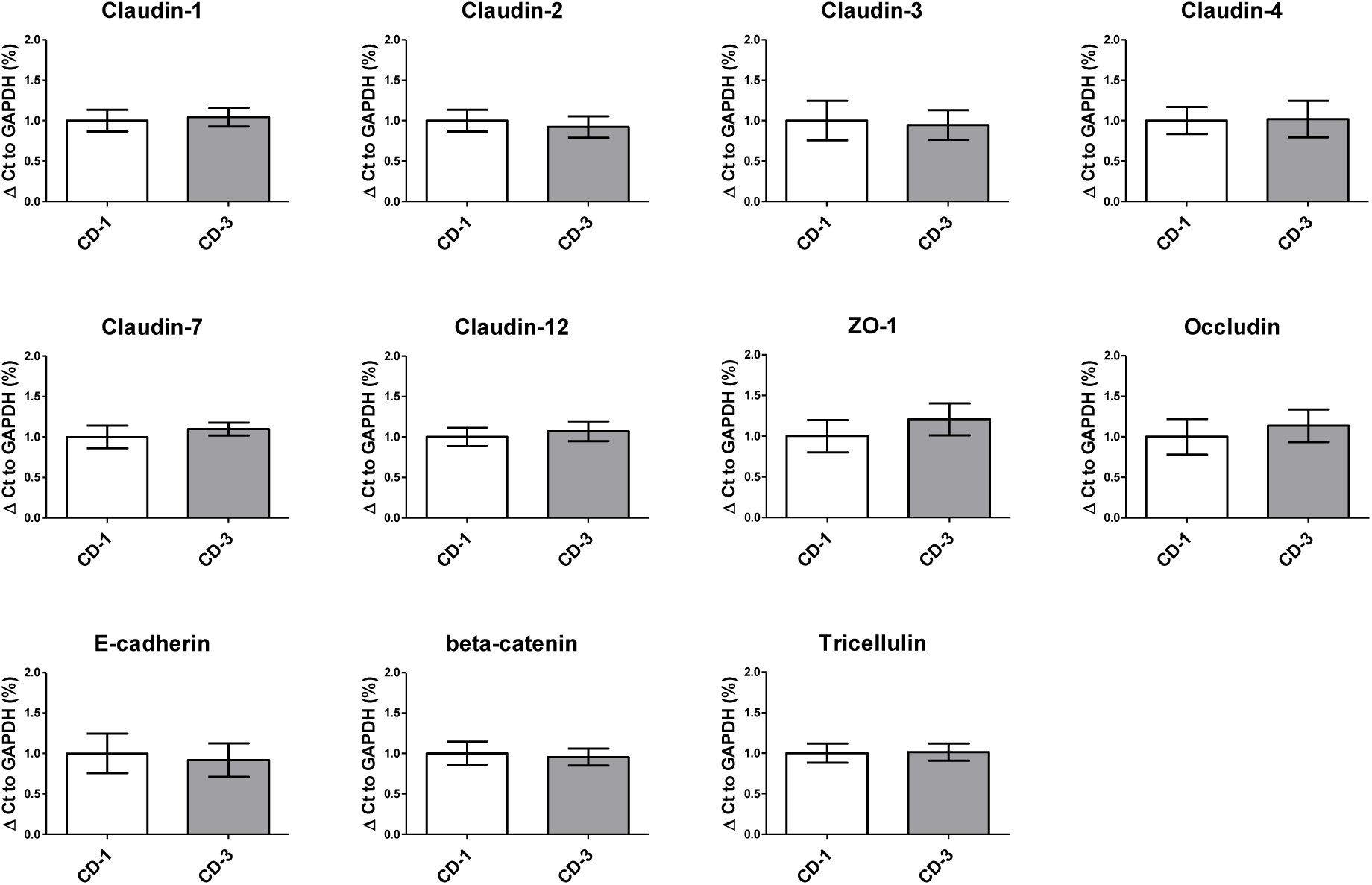
Paracellular junction gene expression in Caco-2 cells exposed to *B. fragilis* culture supernatants remains unaltered. Tight junction (Claudins, ZO-1, Occludin), adherens junction (E-cadherin, beta-catenin) and tricellulin gene expression were unaltered in *B. fragilis bft*-positive CD-3 compared to *bft*-negative CD-1 culture supernatant. Means +/-SD were based on two independent experiments. ZO-1 = Zonulin 1

### Identification of *B. fragilis* genome and metabolome

To identify potential bacterial proteins that may strengthen the epithelial barrier, coding sequences of TEER-elevating strains were compared to those of non-TEER-elevating strains. Herein, nine proteins were identified which were only present in TEER-elevating strains: Bft1, Metalloprotease II (MPII), putative amidoligase, transposase like protein, putative transposase/insertion sequence protein, and four hypothetical proteins.

To further identify metabolites produced by the different *B. fragilis* strains that might contribute to the observed barrier enhancing effect, the concentrated culture supernatants were analyzed using ^1^H Nuclear Magnetic Resonance (NMR) spectroscopy. All strains show comparable metabolic profiles (figure 5A). Based on partial least squares discriminant analysis (PLS-DA) analyses, the TEER-enhancing and non-TEER-enhancing groups could be discriminated based on a decreased relative concentration of acetate and lactate in the TEER-enhancing group (figure 5B).

**Figure 5.**
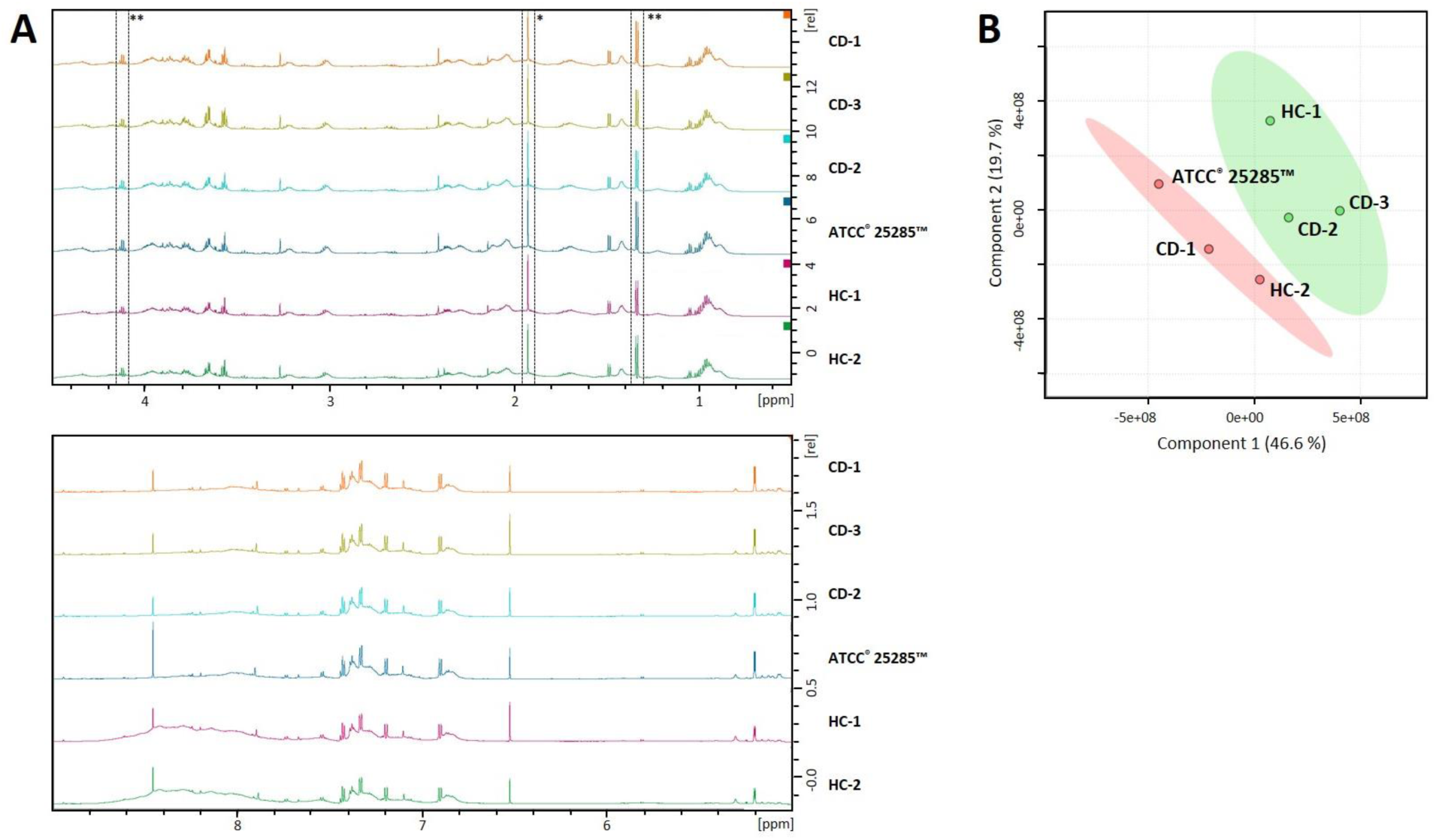
NMR profiles of *B. fragilis* supernatant can distinguish between TEER-enhancing and non-TEER-enhancing strains. (A) The NMR spectra of all strains are largely comparable. Acetate (*) and lactate (**) is relatively decreased in the TEER-enhancing *B. fragilis* strains HC-1, CD-1 and CD-3 compared to all other strains. (B) PLS-DA plot demonstrates that the TEER-enhancing group (green) can be clearly discriminated from the non-TEER-enhancing (red) group.

## Discussion

Previously, *B. fragilis* colonization has been investigated in CD exacerbations, together providing inconclusive results.^4,17,19,25^ Based on previous research, we hypothesized that *B. fragilis* and its virulence factors *bft* and *ubb* are associated with exacerbations and may contribute to exacerbations by disruption of the intestinal epithelial barrier.^26–30^ In the present study, we indeed observed a higher *B. fragilis* prevalence among active CD patients when compared to patients in remission, but not in virulence factor-positive strains. Furthermore, disruption of the intestinal epithelial barrier by supernatant of various (non)virulent *B. fragilis* strains could not be observed. Intriguingly, Bft-positive culture supernatant even led to a significant increase of intestinal epithelial barrier resistance in Caco-2 monolayers.

The cross-sectional investigation of *B. fragilis* in our large CD cohort demonstrated a higher prevalence in active CD patients. However, no differences in relative abundance were detected when analyzing *B. fragilis* positive samples. Logistic regression analysis confirmed the association between *B. fragilis* colonization and disease activity and additionally highlighted an increased likelihood of *B. fragilis* carriage among patients with a stricturing disease behavior. To our knowledge, an association with disease behavior has previously not been reported. Previous research has shown that *B. fragilis* lipopolysaccharide can activate Toll-like receptor 4, which in turn has been associated with liver fibrosis and was suggested as a potential mechanism in stricturing CD.^31,32^ This potential mechanism needs further investigation to clarify whether *B. fragilis* can induce fibrosis and may thereby directly contribute to the development or persistence of strictures.

To study the relevance of *B. fragilis* virulence factors, we next compared the prevalence and relative abundance of *bft* and *ubb* positive strains. Bft and Ubb are two *B. fragilis* proteins, which have previously been shown to affect the host via direct and indirect interactions, respectively.^21,27^ The metalloprotease Bft has been shown to disrupt the epithelial barrier by cleaving the adherens junctions protein E-cadherin *in vitro*,^27^ whereas Ubb was found to mimic human Ubb and led to antibody formation *in vivo*.^21^ These two compounds were therefore considered of interest and detected in the fecal metagenomes of CD patients.

In our cohort, no difference was found in virulence factor positive *B. fragilis* strains when comparing active and remissive CD. Further analyses on associations between virulence factor carriage and CD related parameters were not conclusive due to the overall low prevalence. Although previous rather small scale studies on *bft* in CD patients could also not demonstrate a link with disease exacerbations^25,33^ and studies on *ubb* among CD patients are lacking, their gene products might still play a role in intestinal barrier dysfunction and thereby contribute to intestinal inflammation. We therefore subsequently investigated representative bacterial strains for their effect on intestinal epithelial barrier function using a validated Caco-2 barrier function model.^34^ To investigate the functional impact, we chose to apply bacterial culture supernatant as it contains all secreted products, including proteins, metabolites and MVs.^8–10^ *In vivo*, these products can interact with the host by directly targeting the epithelial lining or indirectly via the immune system.^8,9,35^

In contrast to our hypothesis, none of the investigated supernatants led to a disrupted epithelial barrier when examined with TEER analysis. On the contrary, Bft-positive culture supernatant significantly increased epithelial barrier resistance. Even in the presence of the inflammatory cytokines TNF-α and IFN-γ, which are known to disrupt tight junctions,^18,36^ Bft-positive culture supernatant evoked an equivalent TEER increase. As previous studies have described a barrier disrupting effect induced by Bft in some other colonic (adeno)carcinoma cell lines, such as HT-29 or T84,^26,28^ but not in others, for instance Caco-2, NCI-H508 and LS174T,^37^ we also tested the effect of Bft-positive culture supernatant on a more physiological CD patient-derived colonic organoid model of epithelial barrier function.^38^ This confirmed that exposure to Bft-positive and Bft-negative *B. fragilis* culture supernatant did not lead to barrier disruption or any conspicuous morphological changes, such as cell rounding.

To further elucidate mechanisms underlying the pronounced TEER elevation, junctional gene expression was examined. The qPCR-based approach showed that altered gene expression levels of junctional proteins are unlikely to contribute to the TEER elevation evoked by Bft-positive culture supernatant. Other cellular mechanisms might rather be involved in the detected TEER increase, such as alterations in ion flux or post-translational modifications of junctional proteins.^34,39^ We further excluded a medium-based effect, since the bacterial culture medium was the same for all strains and was also used in the negative control.

Altogether, the above findings are not in line with previous research on the mechanisms of Bft. Several studies showed that isolated Bft could significantly decrease TEER via the cleavage of the adherens junction protein E-cadherin.^26–29^ In those studies, Bft was purified and concentrated using several liters of culture supernatant to reach high toxin concentrations.^26–29,37^ In the present study, we chose to apply the complete culture supernatant of 24 h bacterial cultures instead of purified and enriched Bft, to mimic a more physiological situation. Although we succeeded to detect secreted Bft in the supernatant, using a previously established HT-29 cell rounding assay,^40^ Bft concentrations were below the detection limit of western blot. Furthermore, for barrier function analysis Caco-2 cells were used instead of HT-29 or T84 cells, which might vary in Bft susceptibility.^26,28,37^

However, to confirm that the unexpected lack of epithelial barrier disruption is not merely the result of a potentially more resistant Caco-2 phenotype, we showed that *bft-*positive culture supernatant also did not disrupt the epithelial barrier in more physiological, human colonic organoids as mentioned above. Therefore, we suggest that more physiological concentrations of secreted Bft may act differently on the epithelial barrier than high concentrations.

Based on the previously reported E-cadherin cleavage and subsequent barrier disruption,^26,27^ it seems rather unlikely that Bft is contributing to the observed increased TEER values in Caco-2 monolayers. Therefore, further research was conducted on identifying the potential factor that might induce the observed TEER elevation. As part of the bacterial secretome, *B. fragilis* MVs were isolated from the supernatant and applied on the Caco-2 monolayers. Since TEER values did not change after 24 h incubation, MVs were excluded as TEER elevating factors. Next, whole genome sequencing was applied to identify strain specific coding sequences, indicating differences in secreted proteins, such as sequences related to *bft* and the transposon machinery. Considering their related functions, these proteins seem rather unlikely to induce TEER elevation. However, the close phylogeny of one TEER-elevating and one non-TEER-elevating strain offers limited possibilities for the involvement of other genes. Finally, relative concentrations of acetate and lactate were lower in TEER-increasing strains. However, short chain fatty acids, including acetate, and lactate are reported to rather strengthen the epithelial barrier.^13,41^ The lower acetate and lactate concentrations found in the supernatant of TEER-enhancing strains are therefore unlikely to explain the TEER-enhancing effect. Altogether, we could not elucidate the mechanism underlying the significant increase in TEER by *bft*-positive *B. fragilis* strains.

In summary, this study shows a higher *B. fragilis*, but not *bft* or *ubb*, prevalence in CD exacerbations and an association with a stricturing disease course. Surprisingly, the direct effect of *B. fragilis* products on colonic epithelial cells led to a significant TEER increase in *bft-* positive strains, which might indicate a barrier stabilizing effect. More detailed pathophysiological mechanisms and a potential clinical relevance need further investigation. Additionally, this study clearly stresses the need to investigate functional host-microbe interactions to pursue on taxonomic and functional associations based on microbiome research.

## Patients and Methods

### Patient inclusion

Fecal samples of CD patients participating in the IBDSL biobank project^23^ were available to evaluate the colonization of *B. fragilis* and relevant strains in CD patients and the relation with disease activity. Baseline characteristics, including medication use and demographic information, were extracted from the IBDSL database.^23^

Patients with a fecal calprotectin level > 200 µg/g were defined as active (n = 88) and patients with a fecal calprotectin level < 100 µg/g as remissive (n = 93). Remissive patients without previously reported calprotectin level > 200 µg/g were excluded to avoid misclassification unless the previously elevated calprotectin level was combined with a CRP value < 10 mg/l. In addition, from two patients having a colonoscopy scheduled for clinical reasons, biopsies were collected from the macroscopically non-inflamed tissue of the ascending colon for crypt isolation and subsequent organoid culture.^38^

All patients participated in the IBDSL cohort and gave written informed consent prior to sample collection. The study protocol was approved by the Medical Ethics Committee of the Maastricht University Medical Centre+ (NL31636.068.10) and registered on www.clinicaltrials.gov (NCT02130349).

### Fecal DNA Isolation

Fecal samples stored at −80 °C within 24 h after collection were obtained from the IBDSL biobank^23^ and cut frozen to obtain ∼0.5 g feces. DNA isolation was conducted using Qiagen QIAamp DNA mini kit (Qiagen, ref.: 51306) according to protocol Q of the International Human Microbiome Standards consortium^42^ with minor adjustments in Fastprep(tm) cycles (three series of 1 min of beating at 5.5 ms and 1 min resting) and vacuum drying at 37 °C for 7 min instead of 3 min. Eluted DNA was stored in Buffer AE (Qiagen) at −20 °C until further analysis.

### Enumeration of fecal *B. fragilis*

Metagenomic DNA obtained from the fecal samples was analyzed for the presence and relative abundance of *B. fragilis* as well as the virulence factors Bft and Ubb by means of real-time quantitative PCR. Samples containing *B. fragilis* specific *gyrB* were subsequently analyzed for the presence of *bft* and *ubb*.

The PCR mix for *gyrB* and *bft* contained 5 µl DNA, 12.5 µl Absolute quantitative PCR mix (Abgene, Epsom, UK), 500 mM forward and reverse primer (Sigma-Aldrich; table S1), 250 nM probe and was supplemented with DNAse-free water to a final volume of 25 µl. qPCR and analysis were conducted with 7900HT Fast Real-Time PCR System (Applied Biosystems) and SDS 2.3 software (Applied Biosystems). PCR cycles for *gyrB* and *bft* were 2 min at 50 °C, 10 min at 95 °C, followed by 42 amplification cycles of 15 s at 95 °C and 60 s at 60 °C.

PCR mix for *ubb* and *16S rRNA* contained 2 µl DNA, 12.5 µl Absolute qPCR SYBR Green supermix (Bio-Rad, Hercules, CA), 300 nM forward and reverse primer (Sigma-Aldrich; table S1) and was supplemented with DNAse-free water to a final volume of 25 µl. qPCR and analysis were conducted with MyIQ Single Color Real-Time PCR Detection System (BioRad) and iQ5 software (BioRad). PCR cycles for *ubb* and *16S rRNA* were 3 min at 95 °C, followed by 40, respectively 35 amplification cycles of 95 °C, 20 s at 63 °C, respectively 55 °C, and 30 s at 72 °C. The melting curve was assessed in 60 cycles of 0.5 °C for 10 s each.

### *B. fragilis* isolation from fecal samples

To retrieve toxigenic and non-toxigenic *B. fragilis* strains for *in vitro* experiments, a random selection of fecal samples of three CD patients and two samples of healthy controls (HC) from a previous study that were positive for *B. fragilis* specific *gyrB* were cultured. A portion of ∼1 g was dissolved in Brain Heart Infusion broth (BHI; Sigma Millipore, ref.: 53285, Darmstadt, Germany) containing 4 µg/ml Vancomycin hydrochloride (No 15327, Cayman Chemical, Michigan, USA) and incubated in an anaerobic jar (80 % N_2_, 10 % CO_2_, 20 % H_2_) at 37 °C for 24 h. Next, 1:100 dilutions were inoculated on Bacteroides Bile Esculin Agar with Amikacin (Becton Dickinson, ref.: 254480, Landsmeer, NL) and incubated for 48 h under the same conditions. Medium to large sized single colonies were subsequently inoculated on Columbia Agar with 5 % Sheep Blood (Becton Dickinson, ref.: 254005, Landsmeer, NL) and incubated for 48 h, as described before. This was repeated to guarantee pure cultures. A single colony was then collected in DNAse-free water and checked for *gyrB, bft* and *ubb* using qPCR as described above. The PCR products of *bft* and *ubb* were confirmed using Sanger sequencing to avoid false positive results. To this end, amplicons were purified from remaining nucleotides using the MSB Spin PCRapace-kit (Stratec molecular, ref.: 1020220400) according to manufacturer’s descriptions. 1 µl of DNA was then added to 5.5 µl DNAse-free water, 1 µl forward or reverse primer (2 pmol/l), 1.5 µl BDT 1.1 buffer and 1 µl BDT v1.1 enzyme mix. The DNA was amplified with one cycle of 1 min at 96 °C and 22 cycles of 10 s at 96 °C, 5 s at 58 °C and 2-3 min at 60 °C. Subsequently, the amplicons were sequenced using an ABI 3730 DNA Analyzer (Thermo Fisher Scientific) and were compared to the Nucleotide Basic Local Alignment Search Tool (BLAST; NCBI) database for sequence similarity with previously sequenced *B. fragilis* derived *bft* and *ubb*.

In total, five *B. fragilis* strains were isolated. As no *ubb* positive, *bft-*negative genotype was identified, a reference strain from the American Type Culture Collection (ATCC^®^ 25285(tm)) was obtained. Finally, the following strains were available for further analysis: one *ubb* positive strain (ATCC^®^ 25285(tm)), two *bft-*positive strains (CD-2, CD-3), one strain positive for both *bft* and *ubb* (HC-1) and two strains negative for both virulence factors (HC-2, CD-1; Table S2).

### *B. fragilis* supernatant

To study the impact of excreted *B. fragilis* products on the intestinal barrier *in vitro*, bacterial culture supernatant was used. Therefore, three to five colonies of *B. fragilis* strains were inoculated in BHI (Sigma Millipore, ref.: 53285) for 24 h at 37°C in anaerobic jars. Subsequently, bacterial cultures were centrifuged at 4500 x *g* for 15 min at 4 °C and supernatant was filtered through 0,2 µm pore size syringe filters (Pall Life Sciences, Ref: 4652). To increase the sensitivity, filtered supernatant was concentrated 20x using Amicon® Centrifugal Units 10 K (Merck Millipore, ref.: UFC8010) at 4000 x *g* for 20 min. Cell-free supernatant, stored at 4 °C, gave stable results on TEER for more than 2 months.

### *B. fragilis* MVs

To investigate the impact of MVs on intestinal epithelial barrier function we separated the MVs from the *B. fragilis* cell-free supernatant. First, 30 ml of supernatant (see above) was concentrated to ∼ 250 µl during several centrifugation steps at 4000 x *g* at 4 °C using Ultracentrifugal Filters of 100 kDa (Merck Millipore, ref: UFC9100). After collection of the concentrate, the filter was washed with 250 µl PBS to recover the remaining particles. MVs were isolated by Size Exclusion Chromatography using CL-2B Sepharose (GE healthcare, Little Chalfont, UK). As described by Benedikter *et al*.^43^, vesicle-rich fractions were pooled and the concentration was determined by tunable resistive pulse sensing using qNano Gold (Izon Science Ltd., Oxford, UK) and Izon Control Suite software v3.2.

### Bft secretion and activity

To examine the presence of Bft in the cell-free *B. fragilis* supernatant, we isolated the total protein fraction following the protocol of Casterline *et al*.^44^ with minor modifications. One ml of culture supernatant was precipitated with 10 % (v/v) trichloric acid in aceton containing 20 mM 1,4-dithiothreitol (DTT) for 1 h on ice and centrifuged at 15,000 *x g* for 15 min. The pellet was washed twice with aceton, carefully resuspended and centrifuged at full speed for 15 min. The pellet was then recovered in 2x Laemmli buffer, containing 4 % SDS, 10 % 2-mercaptoethanol, 20 % glycerol, 0.004 % bromophenol blue and 0.125 M Tris-HCl, and stored at −20 °C. For subsequent western blot analysis, the samples were further denatured at 95 °C for 10 min and loaded on a 13 % SDS gel according to standard protocol. After separation and wet blotting on an Amersham™ Protran™ 0.2 µm nitrocellulose membrane (GE Healthcare), the membrane was blocked with skim milk in TBST for 1 h at RT. The membrane was then incubated with anti-Bft antibody (Cusabio, ref.: CSB-PA346537LA01BDP) diluted 1:2000 for 36 h at 4 °C, followed by goat-anti-rabbit antibody conjugated with horseradish peroxidase (Dako p0448) diluted 1:2000 for 30 min at RT and developed with 0.1 M Tris pH 7.5, 0.2 mg/ml DAB, 0.01 % NiCl_2_ and 0.03 % H_2_O_2_. Pictures were taken with Mini HD 9 (Uvitec, Cambridge, UK).

Bft activity was confirmed by a slightly adapted HT-29 cell rounding assay.^40,45^ In brief, HT-29 cells were cultured for five days in a 96-well plate at a density of 2000 cells/well with Dul ecco’s Modified Eagle Medium (DMEM; Sigma Aldrich, ref.: D6429) supplemented with 10 % heat-inactivated fetal bovine serum (FBS; Gibco, ref. 10500) and 1 % antibiotic-antimycotic (100x; anti-anti; Gibco, ref.: 15240-062). Cells were then washed thrice with PBS (Gibco, Life Technologies, ref.: 10010-031) and incubated overnight with 20x concentrated *B. fragilis* cell-free supernatant diluted 1:10 in HT-29 culture medium without FBS at 37 °C. Cell rounding was examined by confocal light microscopy (Leica Microsystems GmbH, Mannheim, Germany), images were taken using LAS-AF software (Leica Microsystems) and analyzed using ImageJ.^46^

### Barrier function analysis

Caco-2 cell monolayers (passage number 47-57), as a well-established *in vitro* model for intestinal epithelial barrier function^34^, were seeded on Millicell Hanging Cell Culture Inserts (Merck Millipore, ref.: MCHT24H48) at a density of 100,000 cells/insert in DMEM supplemented with 1 % FBS, 0.1 % non-essential amino acids (Gibco, ref.: 11140050) and 0.1 % anti-anti. Culture medium was refreshed every 2-3 days in both compartments. Monolayers were allowed to differentiate during 14 to 21 days at 37 °C and 5 % CO_2_ and the TEER was evaluated using the EVOM2 Epithelial Volt/Ohm Meter (World Precision Instruments, Sarasota, FL, USA). Mature monolayers (TEER > 600 Ω*cm^2^) were luminally exposed to BHI or bacterial-free concentrated culture supernatant, each diluted 1:10 in Caco-2 culture medium, or to PBS or a final concentration of 5 × 10^7^ MVs in PBS, each diluted 1:2 in Caco-2 culture medium, and incubated for 24 h. To investigate whether *B. fragilis* supernatant is able to prevent cytokine induced barrier disruption, experimental conditions were pre-incubated basally with 100 ng/ml TNF-α (Sigma, ref.: T6674) in com ination with 100 ng/ml IFN-γ (Sigma, ref.: I3265) for 1 h, before *B. fragilis* supernatant was added luminally. After another 24 h of co-incubation, TEER values were assessed and expressed as percentage of the TEER value prior to incubation.

To confirm that relevant findings were not restricted to Caco-2 cells, concentrated *B. fragilis* culture supernatant was also applied to the more physiological model of colonic patient-derived organoids. Therefore, colonic biopsies were collected in cold PBS, washed thrice with 1 % anti-anti in PBS, thrice with 10 mM DTT in PBS and incubated with 2 mM EDTA in PBS for 1 h at 4 °C and 5 rpm. Biopsies were transferred to PBS and crypts were separated by several rounds of mechanical shaking. The pooled crypt supernatant was then supplemented with 5 % FBS and centrifuged at 400 x *g* for 8 min at 4 °C. The pellet was washed twice with 2 ml cold basal medium (DMEM/F12; Gibco, ref.: 12634-010) supplemented with 1 % GlutaMa (tm) (Life Technologies), 1 % Hepes buffer (Life Technologies) and 5 % FBS). After centrifugation at 400 x *g* for 3 min, crypts were plated in GelTre (tm) (Gi co, ref: A1413201) and IntestiCult(tm) Organoid Growth Medium (StemCell Technology GmbH, Germany, ref.: SC-06010) was added 15 min after incubation at 37 °C, 5 % CO_2_. Medium was refreshed every 3 days and passaged when the morphology appeared complex, according to previous descriptions.^47^ In brief, GelTre (tm) with enclosed organoids was solu ilized on ice for 15 min. Culture medium was replaced by ice cold basal medium. Organoids were disrupted mechanically using a 1000P pipette and centrifuged at 150 x *g* for 5 min. The supernatant was replaced y TrypLE(tm) Express (Thermofisher Scientific, ref: 12605010) containing 10 µM y27632 (Tebu-bio BV) and incubated at 37 °C for 2 min. Organoids were then dissociated mechanically using a firepolished Pasteur pipetted, washed and centrifuged twice with basal culture medium at 400 x *g* for 5 min. Finally, the pellet was resuspended with GelTre (tm) and cultured as descri ed above.

Single lumen organoids (passage numbers 10 & 11) were incubated basolaterally with BHI or *B. fragilis* culture supernatant, diluted 1:10 in organoid medium, or organoid medium only at 37 °C for 24 h. Epithelial barrier function was evaluated by 1 mg/ml FITC-d4 (Sigma) added 6-8 h before confocal light microscopy (Leica Microsystems GmbH, Mannheim, Germany). Images were taken using LAS-AF software and processed using ImageJ.

### Analysis of tight- and adherens junction expression

After TEER measurements of Caco-2 monolayers, RNA was isolated using RNeasy Mini Kit (Qiagen) according to manufacturer’s instructions, including DNAse treatment for 15 min. RNA quantity and purity were determined using a Nanodrop spectrophotometer (NanoDrop Technologies, Wilmington, USA). cDNA was synthesized using 75 % RNA template, 20 % iScript reaction mix and 5 % reverse transcriptase (by iScript cDNA Syntgeses Kit BioRad) and the recommended cycle of 5 min 25 °C, 30 min 42 °C and 5 min 85 °C.

Real-time qPCR was performed according to manufacturer’s instructions using SYBR Green Supermix (BioRad, Veenendal, NL, ref.: 1708885) and CFX 96 Real Time System C1000 Touch Thermal Cycler (Bio-Rad). Primer sequences are listed in table S3. Expression of mRNA was analyzed relative to GAPDH, using the 2^-ΔCt^ method.^48^

### Whole genome sequencing

To characterize the isolated *B. fragilis* strains and to predict possible differences in secreted proteins, the genome of all six strains was sequenced. Genomic DNA was isolated from single colonies resuspended in PBS using MasterPure(tm) Complete DNA and RNA purification kit (Epicenter, MC 85200), following the manufacturer’s protocol. Li rary preparation sequencing and *de novo* assembly was performed as described previously.^49^ Coding sequences were annotated using Prokka (v1.7).^50^ Gene presence/absence was determined using Roary (v3.11.2).^51^ SNP distances among isolates was determined using SKA (v1.0).^52^ All bioinformatics tools were run in default settings.

### Metabolite identification using NMR spectroscopy

To further examine potential differences in excreted metabolites by the different *B. fragilis* strains, the culture supernatants were analyzed using proton (^1^H) NMR spectroscopy. Therefore, 150 µl of 20x concentrated *B. fragilis* culture supernatant was diluted in 420 µl 100 mM phosphate buffer (pH 7.4 at 25 °C) and 30 µl Deuterium oxide (D_2_O, 99 atom %D, 100G, Sigma-Aldrich, Germany) containing 1 mM 3-(trimethylsilyl)propiomic-2,2,3,3-d4 acid (TSP, 98 atom %D TSP, 1G, Sigma-Aldrich, Germany). The samples were then transferred into a 5 mm NMR tube (Bruker samplejet, Sigma-Aldrich, Germany). All ^1^H NMR spectra were recorded on a 700 MHz Bruker Avance spectrometer (Bruker, Germany) at 300 K. Spectra were acquired using a one-dimensional NOESY-presat pulse sequence (RD-90°-t-90°-tm-90°-ACQ), an acquisition time of 2 s, a relaxation delay (D1) of 5 s, mixing time (D8) of 100 ms, receiver gain of 32, 64 scans, 45 K data points and a spectral width of 11161 Hz (15.934 ppm). Spectral preprocessing and preliminary comparisons of the spectra were performed using the Bruker TopSpin 3.2 software.

### Statistical analyses

In order to compare the prevalence and relative abundance of *B. fragilis* (virulence factors) in active and remissive CD patients’ samples, the Χ^2^-test with Yate’s correction and Wilco son rank sum test (for samples tested positive for the respective gene) were performed. To predict carriage of *B. fragilis*, based on the combination of patient baseline characteristics (age, gender, Montreal classification, and medication use), multivariate logistic regression was conducted using backward stepwise regression based on the likelihood ratio. Analyses were computed in IBM SPSS Statistics 25 and statistical significance was considered when *p* < 0.05. Comparing the effects of *B. fragilis* supernatant and MVs on TEER of Caco-2 epithelial monolayers one-way ANOVA and Turkey’s post-hoc test was performed using GraphPad Prism 5. For su sequent analysis of differences in tight junction e pression, student’s t-test was applied. Statistical significance was considered when *p* < 0.05. Analyses were based on at least duplicate experiments with each 3 technical replicates.

Partial Least Squares Discriminant Analysis (PLS-DA) was conducted to identify biologically relevant metabolites that differ between TEER-increasing and non-TEER-increasing groups using MetaboAnalyst v2.0.^53^ The selection of the most discriminating metabolites, that are responsible for the pattern of the PLS-DA score plot, was based on both the interpretation of PLS-loading plot and Variable Importance in Projection (VIP) scores. Metabolites were identified using HMDB 4.0 (The Human Metabolome Database).^54^

## Acknowledgments

The authors would like to thank dr. Annemarie Boleij for sharing her experience, and her support with regard to cell rounding experiments.

## Supplementary tables

**Table S1.**
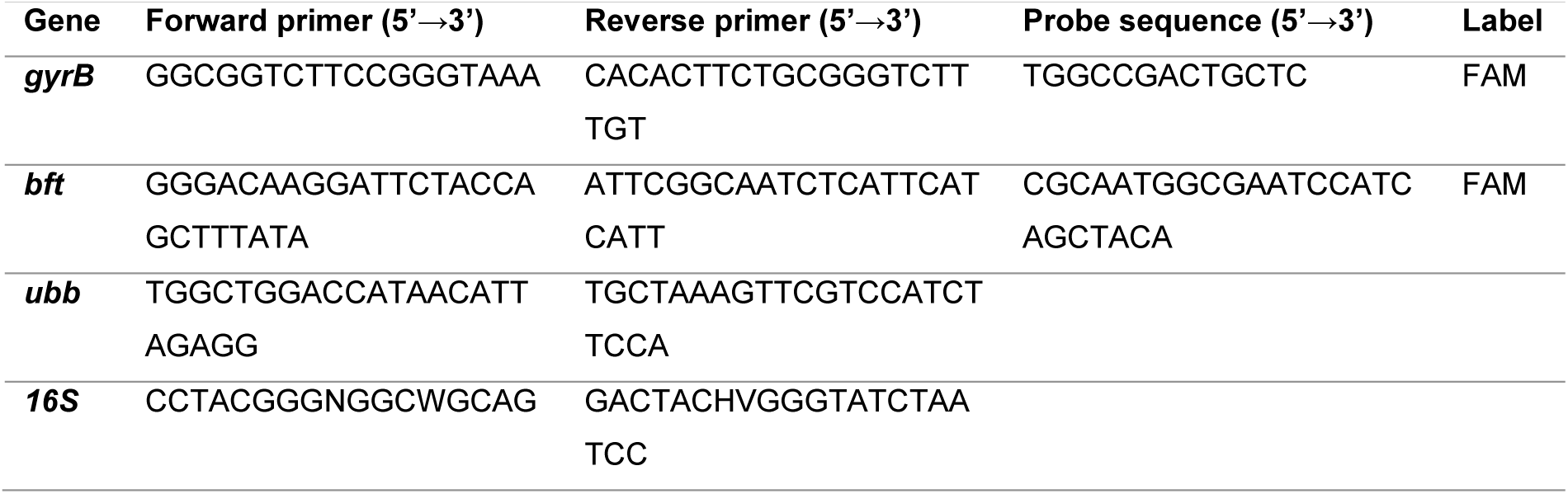
Primer sequences for analysis of prevalence and relative abundance.

**Table S2.** Sample origins of isolated *B. fragilis* strains.

| Subject/Origin | Year | Isolate | Disease status | Calprotectin | Gender | Age |
| --- | --- | --- | --- | --- | --- | --- |
| HC-1 | 2015 | *bft+* *ubb+* | NA | NA | Female | 22 |
| HC-2 | 2015 | *bft-* *ubb-* | NA | NA | Male | NA |
| ATCC® 25285™ | NA | *bft-* *ubb+* | NA | NA | NA | NA |
| CD-1 | 2018 | *bft-* *ubb-* | remission | 59 µg/g | Female | 41 |
| CD-2 | 2018 | *bft+* *ubb-* | remission | 62 µg/g | Female | 29 |
| CD-3 | 2018 | *bft+* *ubb-* | active | 649 µg/g | Male | 50 |

HC = healthy control, NA = not applicable, *bft* = *B. fragilis* toxin, *ubb* = ubiquitin

**Table S3.**
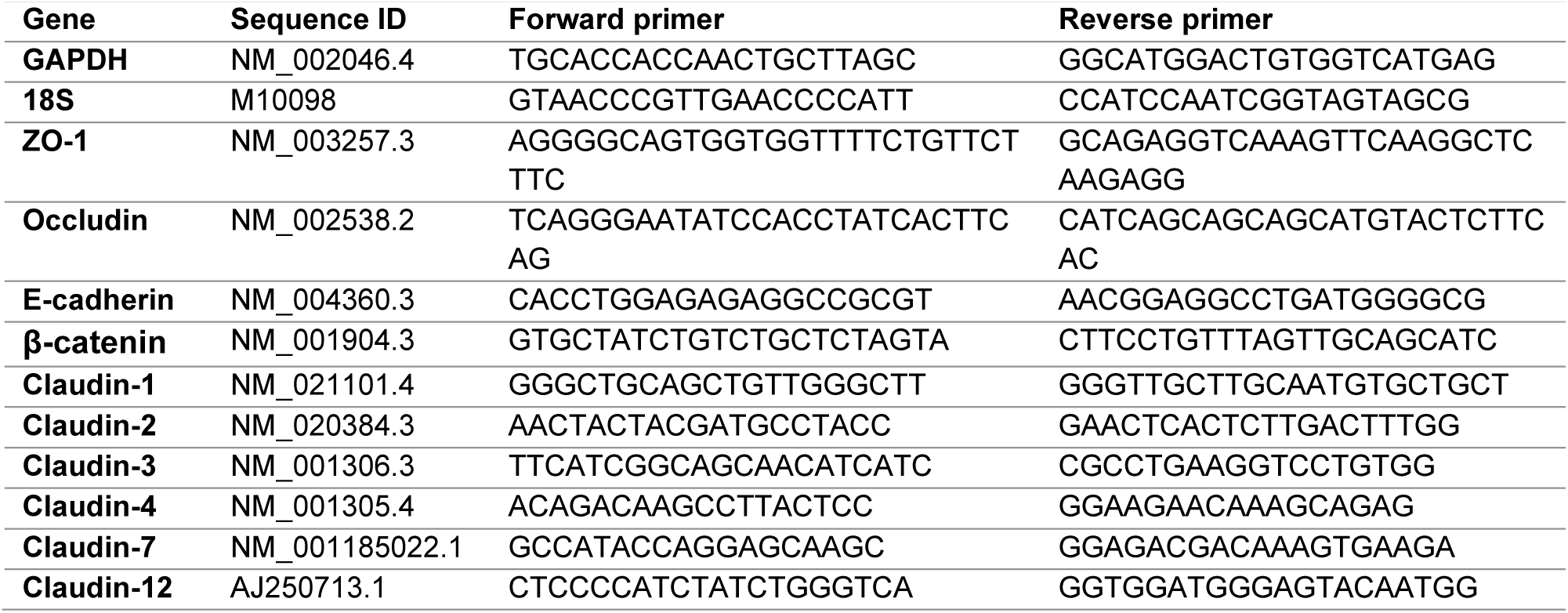
Primer sequences for intercellular junction analysis.

## References

1. Baumgart DC, Sandborn WJ. Crohn’s disease. Lancet. 2012;380:1590–1605.

2. Sol erg IC, Vatn MH, Høie O, et al. Clinical Course in Crohn’s Disease: Results of a Norwegian Population-Based Ten-Year Follow-Up Study. Clin Gastroenterol Hepatol. 2007;5(12):1430–1438. doi: 10.1016/j.cgh.2007.09.002

3. Floyd DN, Langham S, Séverac HC, Levesque BG. The Economic and Quality-of-Life Burden of Crohn’s Disease in Europe and the United States, 2000 to 2013: A Systematic Review. Dig Dis Sci. 2015;60(2):299–312. doi: 10.1007/s10620-014-3368-z

4. Tedjo DI, Smolinska A, Savelkoul PH, et al. The fecal microbiota as a biomarker for disease activity in Crohn’s disease. Sci Rep. 2016;6:1–10. doi: 10.1038/srep35216

5. Galazzo G, Tedjo DI, Wintjens DSJ, et al. Faecal Microbiota Dynamics and their Relation to Disease Course in Crohn’s Disease. J Crohn’s Colitis. 2019;13(10):1273–1282. doi: 10.1093/ecco-jcc/jjz049

6. Prosberg M, Bendtsen F, Vind I, Petersen AM, Gluud LL. The association between the gut microbiota and the inflammatory bowel disease activity: a systematic review and meta-analysis. Scand J Gastroenterol. 2016;51(12):1407–1415. doi: 10.1080/00365521.2016.1216587

7. Li L, Ning Z, Zhang X, et al. RapidAIM: a culture- and metaproteomics-based Rapid Assay of Individual Microbiome responses to drugs. Microbiome. 2020;8(1):1–16. doi: 10.1186/s40168-020-00806-z

8. Kaparakis-Liaskos M, Ferrero RL. Immune modulation by bacterial outer membrane vesicles. Nat Rev Immunol. 2015;15(6):375–387. doi: 10.1038/nri3837

9. Parker A, Lawson MAE, Vaux L, Pin C. Host-microbe interaction in the gastrointestinal tract. Environ Microbiol. 2018;20(7):2337–2353. doi: 10.1111/1462-2920.13926

10. Sicard J-F, Le Bihan G, Vogeleer P, Jacques M, Harel J. Interactions of Intestinal Bacteria with Components of the Intestinal Mucus. Front Cell Infect Microbiol. 2017;7:387. doi: 10.3389/fcimb.2017.00387

11. Caruana JC, Walper SA. Bacterial Membrane Vesicles as Mediators of Microbe – Microbe and Microbe – Host Community Interactions. Front Microbiol. 2020;11(March):1–24. doi: 10.3389/fmicb.2020.00432

12. van Passel MWJ, Kant R, Zoetendal EG, et al. The genome of Akkermansia muciniphila, a dedicated intestinal mucin degrader, and its use in exploring intestinal metagenomes. El-Sayed N, ed. PLoS One. 2011;6(3):e16876. doi: 10.1371/journal.pone.0016876

13. Zheng L, Kelly CJ, Battista KD, et al. Microbial-Derived Butyrate Promotes Epithelial Barrier Function through IL-10 Receptor-Dependent Repression of Claudin-2. J Immunol. 2017;199(8):2976–2984. doi: 10.4049/jimmunol.1700105

14. Sun J, Shen X, Li Y, et al. Therapeutic Potential to Modify the Mucus Barrier in Inflammatory Bowel Disease. Nutrients. 2016;8(1):44. doi: 10.3390/nu8010044

15. Lameris AL, Huybers S, Kaukinen K, et al. Expression profiling of claudins in the human gastrointestinal tract in health and during inflammatory bowel disease. Scand J Gastroenterol. 2013;48:58–69. doi: 10.3109/00365521.2012.741616

16. Zeissig S, Bürgel N, Gu D, et al. Changes in expression and distribution of claudin 2, 5 and 8 lead to discontinuous tight junctions and barrier dysfunction in active Crohn’s disease. Gut. 2007;56:61–72. doi: 10.1136/gut.2006.094375

17. Schäffler H, Kaschitzki A, Alberts C, et al. Alterations in the mucosa-associated acterial composition in Crohn’s disease: a pilot study. Int J Colorectal Dis. 2016;31(5):961–971. doi: 10.1007/s00384-016-2548-z

18. Eeckhaut V, Machiels K, Perrier C, et al. Butyricicoccus pullicaecorum in inflammatory bowel disease. Gut. 2013;62:1745–1752. doi: 10.1136/gutjnl-2012-303611

19. Wills ES, Jonkers DMAE, Savelkoul PH, Masclee AA, Pierik MJ, Penders J. Fecal micro ial composition of ulcerative colitis and Crohn’s disease patients in remission and subsequent exacerbation. PLoS One. 2014;9(3):1–10. doi: 10.1371/journal.pone.0090981

20. Scanlan PD, Shanahan F, Mahony CO, Marchesi JR. Culture-Independent Analyses of Temporal Variation of the Dominant Fecal Microbiota and Targeted Bacterial Su groups in Crohn’s Disease. J Clin Microbiol. 2006;44(11):3980–3988. doi: 10.1128/JCM.00312-06

21. Stewart L, D MEdgar J, Blakely G, Patrick S. Antigenic mimicry of ubiquitin by the gut bacterium Bacteroides fragilis: a potential link with autoimmune disease. Clin Exp Immunol. 2018;194(2):153–165. doi: 10.1111/cei.13195

22. Sears CL. Enterotoxigenic Bacteroides fragilis: a Rogue among Symbiotes. Clin Microbiol Rev. 2009;22(2):349–369. doi: 10.1128/CMR.00053-08

23. van den Heuvel T, Jonkers D, Jeuring S, et al. Cohort Profile: The Inflammatory Bowel Disease South Limburg Cohort (IBDSL). Int J Epidemiol. 2015:1–9. doi: 10.1093/ije/dyv088

24. Brown L, Wolf JM, Prados-Rosales R, Casadevall A. Through the wall: extracellular vesicles in Gram-positive bacteria, mycobacteria and fungi. Nat Rev Microbiol. 2015;13(10):620–630. doi: 10.1016/j.physbeh.2017.03.040

25. Prindiville TP, Sheikh RA, Cohen SH, et al. Bacteroides fragilis enterotoxin gene sequences in patients with inflammatory bowel disease. Emerg Infect Dis. 2000;6(2):171. doi: 10.3201/EID0602.000210

26. Obiso RJ, Azghani AO, Wilkins TD. The Bacteroides fragilis toxin fragilysin disrupts the paracellular barrier of epithelial cells. Infect Immun. 1997;65(4):1431–1439. http://www.ncbi.nlm.nih.gov/pubmed/9119484. Accessed May 6, 2019.

27. Wu S, Lim KC, Huang J, Saidi RF, Sears CL. Bacteroides fragilis enterotoxin cleaves the zonula adherens protein, E-cadherin. Proc Natl Acad Sci. 1998;95(25):14979–14984. doi: 10.1073/pnas.95.25.14979

28. Chambers FG, Koshy SS, Saidi RF, Clark DP, Moore RD, Sears CL. Bacteroides fragilis toxin exhibits polar activity on monolayers of human intestinal epithelial cells (T84 cells) in vitro. Infect Immun. 1997;65(9):3561–3570.

29. Wells CL, Van de Westerlo EMA, Jechorek RP, Feltis BA, Wilkins TD, Erlandsen SL. Bacteroides fragilis enterotoxin modulates epithelial permeability and bacterial internalization by HT-29 enterocytes. Gastroenterology. 1996;110(5):1429–1437. doi: 10.1053/gast.1996.v110.pm8613048

30. Riegler M, Lotz M, Sears C, et al. Bacteroides fragilis toxin 2 damages human colonic mucosa in vitro. Gut. 1999;44(4):504–510. doi: 10.1136/gut.44.4.504

31. Mancuso G, Midiri A, Biondo C, et al. Bacteroides fragilis-derived lipopolysaccharide produces cell activation and lethal toxicity via toll-like receptor 4. Infect Immun. 2005;73(9):5620–5627. doi: 10.1128/IAI.73.9.5620-5627.2005

32. Li J, Mao R, Kurada S, et al. Pathogenesis of fi rostenosing Crohn’s disease. Transl Res. 2019;209:39–54. doi: 10.1016/j.trsl.2019.03.005

33. Zitomersky NL, Atkinson BJ, Franklin SW, et al. Characterization of Adherent Bacteroidales from Intestinal Biopsies of Children and Young Adults with Inflammatory Bowel Disease. PLoS One. 2013;8(6):e63686. doi: 10.1371/journal.pone.0063686

34. Srinivasan B, Kolli AR, Esch MB, Abaci HE, Shuler ML, Hickman JJ. TEER measurement techniques for in vitro barrier model systems. J Lab Autom. 2015;20(2):107–126. doi: 10.1177/2211068214561025

35. Badi SA, Khatami S, Irani S, Siadat SD. Induction Effects of Bacteroides fragilis Derived Outer Membrane Vesicles on Toll Like Receptor 2, Toll Like Receptor 4 Genes Expression and Cytokines Concentration in Human Intestinal Epithelial Cells. Cell J. 2019;21(1):57–61. doi: 10.22074/CELLJ.2019.5750

36. Sturgeon C, Fasano A. Zonulin, a regulator of epithelial and endothelial barrier functions, and its involvement in chronic inflammatory diseases. Tissue Barriers. 2016;4(4):e1251384. doi: 10.1080/21688370.2016.1251384

37. Van Tassell RL, Lyerly DM, Wilkins TD. Purification and characterization of an enterotoxin from Bacteroides fragilis. Infect Immun. 1992;60(4):1343–1350. https://www.ncbi.nlm.nih.gov/pmc/articles/PMC257002/. Accessed May 29, 2019.

38. Xu P, Becker H, Elizalde M, Masclee A, Jonkers D. Intestinal organoid culture model is a valuable system to study epithelial barrier function in IBD. Gut. 2018;67(10):1905–1906. doi: 10.1136/gutjnl-2017-315685

39. Shigetomi K, Ikenouchi J. Regulation of the epithelial barrier by post-translational modifications of tight junction membrane proteins. J Biochem. 2018;163(4):265–272. doi: 10.1093/jb/mvx077

40. Weikel CS, Grieco FD, Reuben J, Myers LL, Sack RB. Human colonic epithelial cells, HT29/C1, treated with crude Bacteroides fragilis enterotoxin dramatically alter their morphology. Infect Immun. 1992;60(2):321–327. http://www.ncbi.nlm.nih.gov/pubmed/1730463. Accessed July 9, 2019.

41. Bruininx EMAM, Koninkx JFJG, Binnendijk GP, et al. Effects of prefermented cereals or the end products of fermentation on growth and metabolism of enterocyte-like Caco-2 cells and on intestinal health of restrictedly fed weanling pigs. animal. 2010;4(1):40–51. doi: 10.1017/S175173110999084X

42. Costea PI, Zeller G, Sunagawa S, et al. Towards standards for human fecal sample processing in metagenomic studies. Nat Biotechnol. 2017;35(11):1069. doi: 10.1038/nbt.3960

43. Benedikter BJ, Bouwman FG, Vajen T, et al. Ultrafiltration combined with size exclusion chromatography efficiently isolates extracellular vesicles from cell culture media for compositional and functional studies. Sci Rep. 2017;7(1):15297. doi: 10.1038/s41598-017-15717-7

44. Casterline BW, Hecht AL, Choi VM, Bubeck Wardenburg J. The Bacteroides fragilis pathogenicity island links virulence and strain competition. Gut Microbes. 2017;0976:1–10. doi: 10.1080/19490976.2017.1290758

45. Mundy LM, Sears CL. Detection of Toxin Production by Bacteroides fragilis: Assay Development and Screening of Extraintestinal Clinical Isolates. Clin Infect Dis. 1996;23(2):269–276. doi: 10.1093/clinids/23.2.269

46. Abràmoff MD, Magalhães PJ, Ram SJ. Image processing with imageJ. Biophotonics Int. 2004;11(7):36–41. doi: 10.1201/9781420005615.ax4

47. Sato T, Stange DE, Ferrante M, et al. Long-term Expansion of Epithelial Organoids From Human Colon, Adenoma, Adenocarcinoma, and Barrett’s Epithelium. Gastroenterology. 2011;141(5):1539–1541. doi: 10.1053/j.gastro.2011.07.050

48. Schmittgen TD, Livak KJ. Analyzing real-time PCR data by the comparative CT method. Nat Protoc. 2008;3(6):1101–1108. doi: 10.1038/nprot.2008.73

49. Flipse J, von Wintersdorff CJH, van Niekerk JM, et al. Appearance of vanD-positive Enterococcus faecium in a tertiary hospital in the Netherlands: prevalence of vanC and vanD in hospitalized patients. Sci Rep. 2019;9(1):6949. doi: 10.1038/s41598-019-42824-4

50. Seemann T. Prokka: rapid prokaryotic genome annotation. Bioinformatics. 2014;30(14):2068–2069. doi: 10.1093/bioinformatics/btu153

51. Page AJ, Cummins CA, Hunt M, et al. Roary: rapid large-scale prokaryote pan genome analysis. Bioinformatics. 2015;31(22):3691–3693. doi: 10.1093/bioinformatics/btv421

52. Harris SR. SKA: Split Kmer Analysis Toolkit for Bacterial Genomic Epidemiology. bioRxiv. October 2018:453142. doi: 10.1101/453142

53. Chong J, Yamamoto M, Xia J. MetaboAnalyst 2.0: From Raw Spectra to Biological Insights. Metabolites. 2019;9(3):57. doi: 10.3390/metabo9030057

54. Wishart D, Feunang Y, Macu A, Guo A, Liang K. HMDB 4.0 - The Human Metabolome Database for 2018. Nucleic Acids Res. 2018;46(D1):D608–617. doi: 10.1093/nar/gkx1089

